# GPR27 mediates adrenergic ligands-induced transinhibition of EGFR

**DOI:** 10.1101/2025.03.12.642381

**Authors:** Sorina Andreea Anghel, Rodica Badea, Cosmin Trif, Teodora Stratulat, Cristiana Trita, Diana-Georgiana Navligu, Stefana M. Petrescu, Alexandru Babes, Costin-Ioan Popescu, Cristin Coman, Julien Hanson, Sorin Tunaru

**Affiliations:** Cell Signalling Research Group, Institute of Biochemistry of the Romanian Academy, Splaiul Independentei 296, 060031, Bucharest, Romania; Department of Anatomy, Physiology, and Biophysics, Faculty of Biology, University of Bucharest, Splaiul Independenţei 91-95, 050095 Bucharest, Romania; Institute of Biochemistry of the Romanian Academy, Department of Viral Glycoproteins, 060031 Bucharest, Romania; “Cantacuzino” Medico-Military National Research Institute, 050096 Bucharest, Romania; Laboratory of Molecular Pharmacology, GIGA-Molecular Biology of Diseases, University of Liege, Liege, Belgium; Laboratory of Medicinal Chemistry, Center for Interdisciplinary Research on Medicines (CIRM), University of Liege, Liege, Belgium; Prothanor Biotech S.R.L, Ing. Ion Zablovschi 81, Sector 1, Bucharest, Romania

**Author notes:** Corresponding author: Sorin Tunaru. **Funding:** This work was funded by an EEA-Romania-Norway research grant (2018-0535) entitled “New Generation of Drug Targets for Schizophrenia”, (NEXTDRUG). **Competing Interests:** The authors have no relevant financial or non-financial interests to disclose.

## Abstract

G-protein coupled receptor 27 (GPR27) is part of the “Super Conserved Receptors Expressed in Brain” (SREB) family, alongside GPR85 and GPR173. While the endogenous ligands and functions of SREB receptors are still unknown, GPR27 has been implicated in insulin secretion and tumorigenesis. Here, we show that substituting GPR27’s C-terminus domain with that of the β1 adrenergic receptor (β1AR) yields a chimera with β1AR-like ligand selectivity and cellular functions. Interestingly, adrenergic ligands stimulation of GPR27 inhibited EGF-induced serum-responsive element (SRE) activation, independently of G-proteins and β-arrestins, through dephosphorylation of c-Src and EGFR proteins. This unique response was exclusive to GPR27, as GPR85 and GPR173 showed no similar effects. These findings suggest that GPR27 is a receptor responding to adrenergic ligands to transinhibit EGFR through an atypical signaling mechanism.

## 2. Introduction

G protein-coupled receptors (GPCRs) represent one of the largest and most diverse families of membrane receptors in mammalian cells, playing pivotal roles in transducing extracellular signals into intracellular responses (1). Among the less explored members of this vast superfamily are the Super Conserved Receptors Expressed in the Brain (SREB), which include SREB1 (GPR27), SREB2 (GPR85), and SREB3 (GPR173). These receptors, first identified through genomic analysis and named for their conserved sequences across species, are predominantly expressed in the central nervous system (CNS), hinting at their potential roles in neurophysiology and brain function (2, 3). Despite their evolutionary conservation and high expression in CNS, the endogenous ligands and physiological functions of SREBs have remained elusive, classifying them as orphan GPCRs (4). Recent research, however, has begun to shed light on the potential roles of these receptors. GPR27 for example has been implicated in modulating insulin secretion, suggesting a link between GPR27 and metabolic regulation (5, 6). Moreover, the heterologous expression of GPR27 in HEK293T cells resulted in elevated inositol phosphate levels pointing to a Gq/11 coupling of this receptor (7). On the other hand, a broader study involving the analysis of the transient expression of a large number of GPCRs on cAMP levels in Chinese hamster ovary (CHO) cells revealed that GPR27, but not GPR85 and GPR173, display significant constitutive inhibitory activity on cAMP-driven gene expression, suggesting potential Gi-proteins coupling of GPR27 (8). In addition, several synthetic agonists of GPR27 have been recently proposed. These agonists display specificity for GPR27 over GPR85 and GPR173 and, interestingly, in the case of GR27, induced the recruitment of β-arrestin 2 in a G-protein-independent manner (9). Evidence of possible implications in pathological conditions was drawn from the analysis of The Cancer Genome Atlas Database, where lower expression levels of GPR27 were correlated with higher death rates in patients with glioma (10).

The functional interaction between GPCRs and receptor tyrosine kinases (RTKs) has been the subject of intense research since the seminal work by Ulrich and colleagues describing tyrosine phosphorylation of the epidermal growth factor receptor (EGFR) in response to known GPCR agonists, a phenomenon termed “transactivation” (11). EGFR is not the only RTK family member subject to transactivation by GPCRs. Other RTKs such as fibroblast growth factor receptors (FGFRs), platelet-derived growth factor receptor (PDGFR), or the insulin receptor (IR) have been reported to be targets for the GPCRs transactivation (12). The biological consequences of transactivation are well described, for example, the activation of GPCRs by ligands such as lysophosphatidic acid (LPA), endothelin-1, and thrombin in Rat-1 fibroblasts promotes S phase entry and DNA synthesis through transactivation of the EGFR (13). The molecular mechanisms underlying this phenomenon are still not fully clear. Several lines of evidence demonstrate that the transactivation of RTKs by GPCRs involves metalloproteinases (MMPs) or disintegrin and metalloproteinases (ADAMs). MMPs and ADAMs cleave pro-ligands of RTKs bound to the extracellular matrix components leading to the release of the ligands that can bind to their cognate RTKs triggering intracellular signaling events (14, 15). In contrast to the transactivation phenomenon, the inhibition of RTKs signaling by GPCRs has not yet been widely reported. However, CXCL12 stimulation of C-X-C chemokine receptor type 4 (CXCR-4) positive HeLa and 5637 cells induced the inhibition of EGFR phosphorylation involving protein kinase C (PKC) (16). Although the transactivation of RTKs by GPCRs has been associated with important biological functions, the role of transinhibition has not yet been described. The present study shows that GPR27 is a novel adrenergic receptor functioning atypically to transinhibit RTKs such as EGFR without engaging G-proteins and arrestins.

## 3. Materials and Methods

### 3.1 Chemicals

The majority of chemicals were purchased from Cayman Chemicals, with a few exceptions. Pertussis toxin (PTX) was obtained from Invitrogen™ (Carlsbad, CA, USA), YM-254890 from Focus Biomolecules (Montgomery County, PA, USA), the GRK inhibitor, 4-Amino-5-(bromomethyl)-2-methyl pyrimidine dihydrobromide (CAS 5423-98-3) from Santa Cruz Biotechnology (Texas, USA), polyethyleneimine (PEI), gallein, PP2 (4-amino-5-(4-chlorophenyl)-7-(t-butyl)pyrazolo[3,4-d]pyramidine) were from Sigma-Aldrich (St. Louis, MO, USA) and the epidermal growth factor (EGF) was obtained from PeproTech Inc. (Rocky Hill, New Jersey, USA). The FDA-Approved Drug Library containing 1403 compounds was acquired from TargetMol (Boston, MA, USA).

### 3.2 Cell culture and transfection

HEK293T cells (RRID: CVCL_0063) were grown in DMEM with 4.5 g/L glucose (Pan Biotech, Aidenbach, Germany) supplemented with 10% foetal bovine serum (FBS) (Gibco, Thermo Fisher Scientific, Waltham, MA, USA) and 1% penicillin and streptomycin mix (Gibco) at 37°C and 5% CO2. Plasmid constructs were transfected in HEK-293T cells using PEI.

### 3.3 Plasmids

Human GPR27 (SREB1) and GPR173 (SREB3) cDNAs were cloned into pcDNA3.1-N-DYK vector (GenScript Biotech Corporation, Nanjing, China). Human GPR85 (SREB2) was amplified by PCR using pcDNA3.1(+)-GPR85 (cDNA Resource Center, The Bloomsburg University Foundation, PA, USA) as template and the following primers: forward - 5’-CCTAGGTACCATGGCGAACTATAGCC - 3’, reverse - 5’ – GCTCGGATCCTCATATAACACAGTAAGG - 3’

The amplified fragment was inserted into pcDNA3.1-N-DYK vector between KpnI and BamHI restriction sites. pGloSensorTM-22F cAMP plasmid, pGL4.29[luc2P/CRE/Hygro], pGL4.33[luc2P/SRE/Hygro] were purchased from Promega Corporation (Madison, WI, USA). pOZITX-S1 was a gift from Jonathan Javitch (Addgene plasmid #184925; http://n2t.net/addgene:184925; RRID: Addgene_184925). N-terminally HA-tagged β1AR and the chimeric receptors cDNAs were synthesized and cloned into pcDNA3.1(+). The truncated form of GPR27 (GPR27ΔCterm) was generated by PCR cloning using pcDNA3.1-N-FLAG-GPR27 as a template and the following primers: forward - 5’-AATGGATCCACCATGAAGACGATCATC-3’, reverse - 5’-TAGTCTAGATTAGCTCTGGCAGCAGGGGAAC - 3’.

The amplicon product was then inserted into pcDNA3.1(+) vector between BamHI and XbaI restriction sites. pIRES-N-HA-FLuc-Arrb2 encoding the rat β-arrestin 2, pcDNA3.1-N-FLAG-GPR27-FLuc, pcDNA3.1-N-FLAG-β2AR-FLuc, pcDNA3.1-N-FLAG-GPR27V2-Fluc and pcDNA3.1-N-FLAG-β2ARV2-Fluc. The GPR27 mutant Y224F was constructed using the Q5® Site-Directed Mutagenesis Kit (New England Biolabs, Ipswich, MA, USA) starting from pcDNA3.1-N-FLAG-GPR27 with the primer pair:

5’-ACCTCGTCTTCCTCCGCCTG-3’,

5’-GCGTGGCGCCCACCACCA-3’.

PWPXLD-EGFR-WT was a gift from Chay Kuo (Addgene plasmid #133749; http://n2t.net/addgene:133749; RRID: Addgene_133749). All plasmids generated were validated by enzymatic digest and sequencing (CeMIA, Larissa, Greece).

### 3.4 Determination of cell surface receptor expression by whole-cell ELISA

HEK293T cells were transiently transfected with plasmids encoding receptors cDNAs in a 96-well plate format. The following day, cells were washed with Hanks’ Balanced Salt Solution (HBSS) and fixed with paraformaldehyde 1% for 5 minutes. The permeabilization of cells was performed using TritonX-100 0.1% treatment for 3 minutes to assess the total protein expression. Then, cells were washed three times with HBSS, blocked with 5% non-fat dry milk/HBSS for 1h, and probed with horseradish peroxidase-conjugated antibody directed against the FLAG epitope (1:1000, Sigma) for another hour. After the washing step, cells were incubated with TMB substrate (BD Biosciences, SanDiego, USA), and the reaction was stopped by adding 1M H_2_SO_4_, resulting in a yellow color solution with an absorbance at 450 nm that was measured on a microplate reader (Mithras LB 940, Berthold Technologies).

### 3.5 cAMP determination

HEK293T cells were seeded in white clear bottom 96-well plates and transiently co-transfected with a plasmid encoding a cytosolic localized cAMP-sensitive bioluminescent probe, pGlo-22F (Promega), and plasmids containing receptor cDNAs. After 24h, cells were incubated at room temperature in HBSS containing 1 mM Luciferin EF (NanoLight Technology, AZ, USA) for 2h, in the dark. Following ligands stimulation for 15 minutes, luminescence was recorded by a microplate reader (Mithras LB 940, Berthold Technologies).

### 3.6 GTPase assay

HEK293T cells seeded in 55 cm^2^ dishes were transiently transfected with the pcDNA3.1 empty plasmid and with plasmids encoding GPR27 and β1AR cDNAs, respectively. Twenty-four hours later, cells were washed with cold PBS and lysed with hypotonic buffer (20 mM HEPES, pH 7.4, 1 mM EDTA supplemented with 1 mM DTT, protease inhibitor cocktail EDTA-free (Roche Diagnostics GmbH, Mannheim, Germany), 1 mM PMSF, 1 mM orthovanadate, 5 mM iodoacetic acid, 5 mM NaF) by passing them 15-20 times through a 26G syringe needle. Then, the lysate was centrifuged for 10 min at 9000xg, 4°C and the nuclear fraction was discarded. The soluble fraction was further centrifuged for 40 minutes at 25000xg, 4°C and the resulted pellet (membrane fraction) was resuspended in GTPase/GAP buffer (from GTPase-GloTM Assay kit) containing or not 2 nM isoproterenol. GTPase activity was further determined using the GTPase-GloTM assay kit (Promega) following the protocol for intrinsic GTPase activity recommended by the manufacturer. After adding the Detection Reagent, the reactions were incubated for 5-10 minutes at 25°C and the luminescence was recorded on a microplate reader (Mithras LB 940, Berthold Technologies). Total protein of each sample was estimated using a Pierce BCA Protein Assay kit (Thermo Fisher Scientific, Waltham, MA, USA). To determine the specific GTPase activity, the luminescence value obtained was normalized to the total protein for each sample.

### 3.7 CRE- and SRE-luciferase reporter gene assay

HEK293T cells were seeded in white clear bottom 96-well plates and transiently transfected with pGL4.29[luc2P/CRE/Hygro] or pGL4.33[luc2P/SRE/Hygro] (Promega) together with plasmid encoding receptors cDNAs. After 24h, cells were incubated for 4h at 37°C, 5% CO2 with different compounds in serum-free media and lysed with 0.25% Triton X-100 for 30 minutes at 4°C. Luciferase activity was measured using Firefly Luciferase Assay Reagent (NanoLight) on a microplate reader (Mithras LB 940, Berthold Technologies). For SRE and SRF-RE experiments, media was replaced by DMEM containing 2% FBS before transfection and 1% FBS was used during the assay.

### 3.8 Beta-arrestin 2 recruitment assay

β-Arrestin 2 recruitment assay was performed by Firefly Luciferase Complementation Assay as previously described (9). Briefly, cells were transfected with plasmids encoding cDNAs for β-arrestin 2 fused with the N-terminal part of the firefly luciferase (β-arr2-FLuc) and GPR27 or β2AR fused with the C-terminal part of the firefly luciferase (GPR27-FLuc or β2AR-FLuc). Additionally, we employed chimeric receptors consisting of GPR27 or β2AR with the C-terminus domain replaced by the homologous domain of vasopressin 2 receptor (V2R) fused with the C-terminal part of the firefly luciferase (GPR27-V2R-Fluc or β2AR-V2R-Fluc). After 24h, the medium was replaced with HBSS, and cells were stimulated with compounds for 10 minutes at room temperature. Luminescence was recorded on a microplate reader (Mithras LB 940, Berthold Technologies) after luciferin EF (1 mM) addition.

### 3.9 Immunoblotting

HEK293T cells seeded in 12-well plates were transiently transfected with plasmids containing receptors cDNAs. After 24h, cells were starved for 6h or overnight (16h) in DMEM without FBS and, then treated with isoproterenol at the indicated concentrations, for 10 minutes at 37°C. Then, the cells were lysed in ice-cold radioimmunoprecipitation assay (RIPA) buffer (50 mM Tris, 150 mM NaCl,1% NP-40, 0.5% sodium deoxycholate, 0.1% SDS, pH 7.5) supplemented with protease inhibitors cocktail (Thermo Fisher Scientific, Waltham, MA, USA) and the lysates were clarified by centrifugation 45 minutes at 25000xg, 4°C. Equal amounts of total lysates were separated by SDS-PAGE and transferred on a PVDF membrane. Detection was performed with specific antibodies: rabbit monoclonal anti-pERK (1:1000), rabbit monoclonal anti-ERK (1:1000), rabbit monoclonal anti-pY416 (c-Src) (1:1000), and anti-HA-HRP (1:1000) all from Cell Signaling Technology (Danvers, MA, USA), mouse monoclonal anti-c-Src (Merck-Millipore, Darmstadt, Germany), mouse monoclonal anti-FLAG-HRP (Sigma-Aldrich, Darmstadt, Germany), rabbit polyclonal anti-calnexin (1:6000) (kindly provided by Dr. Gabriela Chiritoiu, Institute of Biochemistry, Bucharest, Romania). As secondary antibodies we used goat anti-mouse-HRP (1:4000) and mouse anti-rabbit-HRP (1:4000).

### 3.10 Statistics

The presented data are expressed as mean ± SEM. Statistical analysis was performed by using GraphPad Prism 6 software. Unpaired Student’s t-test was used to compare the two groups.

## 4. Results

### 4.1 Heterologously expressed human GPR27 reduces intracellular cAMP levels in a Gi/o/z-protein-independent manner

To explore the cellular functions of SREBs, we examined whether the heterologous expression of human GPR27, GPR85, and GPR173 in HEK293T cells impacts various intracellular signaling pathways, including those modulating cAMP levels. As illustrated in Figure 1a, expression of GPR27 and GPR173 significantly reduced forskolin-induced cAMP accumulation, with the most pronounced effect observed for GPR27, whereas GPR85 showed no impact on intracellular cAMP levels. Based on the observation that GPR27 severely affected basal-as well as forskolin-induced increases in cAMP, we next examined the possibility that GPR27 is a constitutively active receptor interacting with Gi-proteins as previously reported (8). To explore this hypothesis, we determined the effect of stimulation with 10 µM forskolin on cAMP levels in HEK293T cells expressing GPR27 in the absence and the presence of the Gi/o proteins inhibitor pertussis toxin (PTX). As presented in Figure 1b, pretreatment of cells with PTX did not result in the reversal of the inhibitory effect of GPR27 on cAMP levels following forskolin stimulation. However, the well-known inhibitory effects of nicotinic acid (niacin) on cAMP mediated by the hydroxycarboxylic acid receptor 2 (HCA_2_R) (17) were fully reversed by PTX pretreatment, confirming the toxin’s effectiveness. To further address the potential involvement of other types of G-proteins, such as Gz, we examined the effect of a recently described toxin, OZITX (Gα<u>o</u>, Gα<u>z,</u> and Gαi inhibiting to<u>x</u>in) (18) on the ability of GPR27 to decrease cAMP levels in cells stimulated with forskolin. As shown in Figure 1c, the expression of OZITX did not reverse the inhibitory effects of GPR27 on cAMP levels increased by forskolin. However, the quinpirole inhibition of forskolin-induced increase in cAMP levels through the dopamine D2 receptor was fully reversed by the presence of OZITX. Although GPR27 showed significant cellular effects in the absence of ligand stimulation which were not mediated by Gi/o/z-proteins, it localized at the plasma membrane as demonstrated by the detection of an anti-FLAG-HRP conjugated antibody in non-permeabilized and permeabilized conditions of HEK293T cells expressing human N-terminally FLAG-tagged GPR27 (Figure 1d). These results demonstrate that GPR27 is not a Gi/o/z-coupled constitutively active receptor, as it strongly inhibits the accumulation of intracellular cAMP independently of these types of G-proteins.

**Figure 1.**
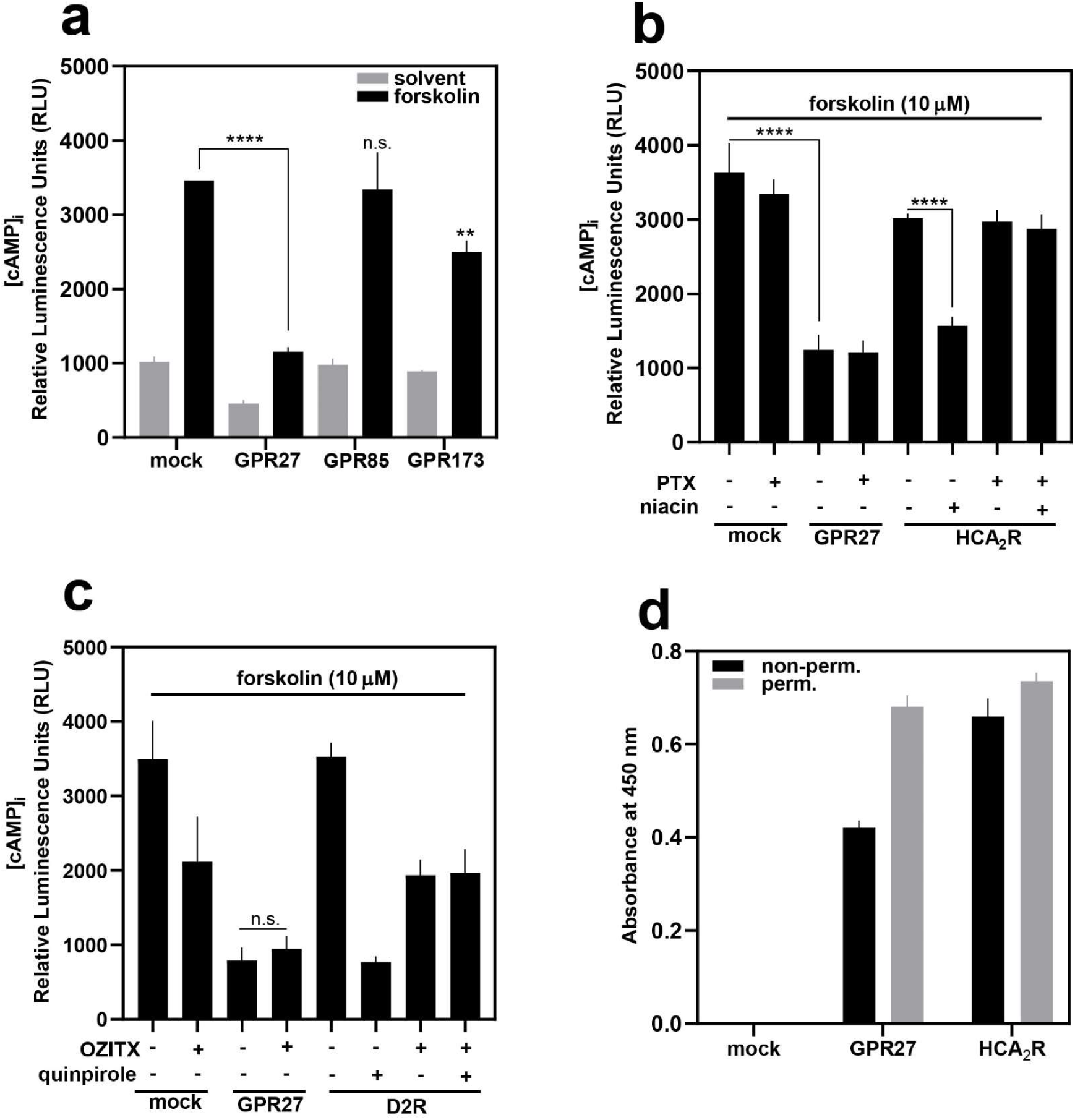
GPR27 inhibits forskolin-induced cAMP increases in a Gi/o/z-independent manner. (a) comparison between the effect of heterologous expression of GPR27, GPR85, GPR173, and empty vector (mock) on basal (solvent)- and 10 µM forskolin-induced cAMP increases in HEK293T cells. (b) effect of pertussis toxin (PTX, 100 ng/ml) pretreatment of cells expressing GPR27, HCA2R, and empty vector on forskolin-induced cAMP increase. Cells expressing the HCA2 receptor were stimulated with 100 µM nicotinic acid (niacin) in the absence or presence of PTX pretreatment to ensure the toxin’s effectiveness. (c) effect of OZITX expression in HEK293T cells with empty vector, GPR27, and D2R on forskolin (10 µM)-induced cAMP increases. D2R agonist quinpirole (10 µM) stimulation of cells co-expressing OZITX or empty vector and D2R was employed to demonstrate the toxin’s effectiveness. (d) plasma membrane and overall cellular expression of N-terminally FLAG-tagged GPR27 in comparison with a typical Gi-coupled receptor, N-terminally FLAG-tagged HCA2R in HEK293T cells. Colorimetric reaction was initiated and absorbance was recorded after exposure of cells to anti-FLAG-HRP-conjugated antibody, followed by preparative steps as described in Material and Methods. Shown are mean values ± SEM, n = 4, *\*\*P* ≤ 0.01 (compared to forskolin-induced cAMP increases in mock-transfected cells), *\*\*\*\*P* ≤ 0.0001, (Student’s two-tailed *t*-test).

### 4.2 Critical role of C-terminus domains of GPR27 and β1AR in signaling

Given that heterologous expression of GPR27 induces cellular effects independently of ligand stimulation and without engaging G-proteins, we investigated whether the intracellular C-terminal domain of GPR27 might be responsible for these effects. To explore this idea, we employed several strategies, including the construction of chimeric GPR27 receptors containing the C-terminal domains of receptors with known ligands. Functional studies were then conducted to evaluate the impact of these modifications on chimera signaling. Additionally, we created chimeras where the C-terminus of known GPCRs was replaced with that of GPR27 and assessed their functional properties (Fig. 2a, d, inside panel). Surprisingly, the chimera of GPR27 containing the β1-adrenergic receptor (β1AR) C-terminus (GPR27-Ctβ1) responded to increasing concentrations of isoproterenol in the cAMP accumulation assay, mimicking β1AR, while wild-type GPR27 remained unresponsive to concentrations up to 30 nM. Conversely, the chimera of β1AR containing the GPR27 C-terminus (β1AR-Ct27) was unresponsive to isoproterenol, similar to GPR27 (Fig. 2b). These functional differences were not due to altered expression levels, as both GPR27-Ctβ1 and β1AR-Ct27 showed comparable plasma membrane localization in HEK293T cells (Suppl. Fig. S1). Similar results were observed in the cAMP accumulation assay when both chimeras were exposed to increasing concentrations of the endogenous β1AR ligand, adrenaline (Fig. 2c). These findings prompted us to assess the impact of FDA-approved compounds on GPR27-Ctβ1 and β1AR. As shown in Table 1, adrenergic and dopaminergic ligands induced similar activation profiles on GPR27-Ctβ1 and β1AR. The observation that the β1AR C-terminus confers GPR27 responsiveness to adrenergic ligands led us to investigate whether C-termini from other adrenergic receptors, such as β2AR or α1AR, would impart similar functional properties to GPR27. As shown in Figure 2d, replacing the GPR27 C-terminus with the corresponding domains of β2AR (GPR27-Ctβ2) or α1AR (GPR27-Ctα1) did not result in chimeras capable of mediating cAMP increases following stimulation with maximal concentrations of isoproterenol. These results indicate that the unique and intriguing combination of GPR27’s seven-transmembrane domain and the β1AR C-terminus is necessary to create a chimeric receptor that mirrors β1AR’s ligand recognition and Gs-mediated signaling. Given the responsiveness of the GPR27-Ctβ1 chimera to adrenergic ligands, we further examined whether wild-type GPR27 mediates isoproterenol effects by stimulating Gs-induced intracellular cAMP increases or by inhibiting forskolin-induced cAMP elevation. Stimulation of HEK293T cells expressing GPR27 with a maximal concentration of isoproterenol did not lead to cAMP accumulation, unlike the robust cAMP increase observed in β1AR-expressing cells (Suppl. Fig. S2a). Additionally, isoproterenol stimulation of GPR27-expressing cells failed to inhibit forskolin-induced cAMP elevation (Suppl. Fig. S2b). These findings demonstrate that GPR27 does not engage Gs- or Gi-proteins to alter intracellular cAMP levels in response to isoproterenol stimulation.

**Figure 2.**
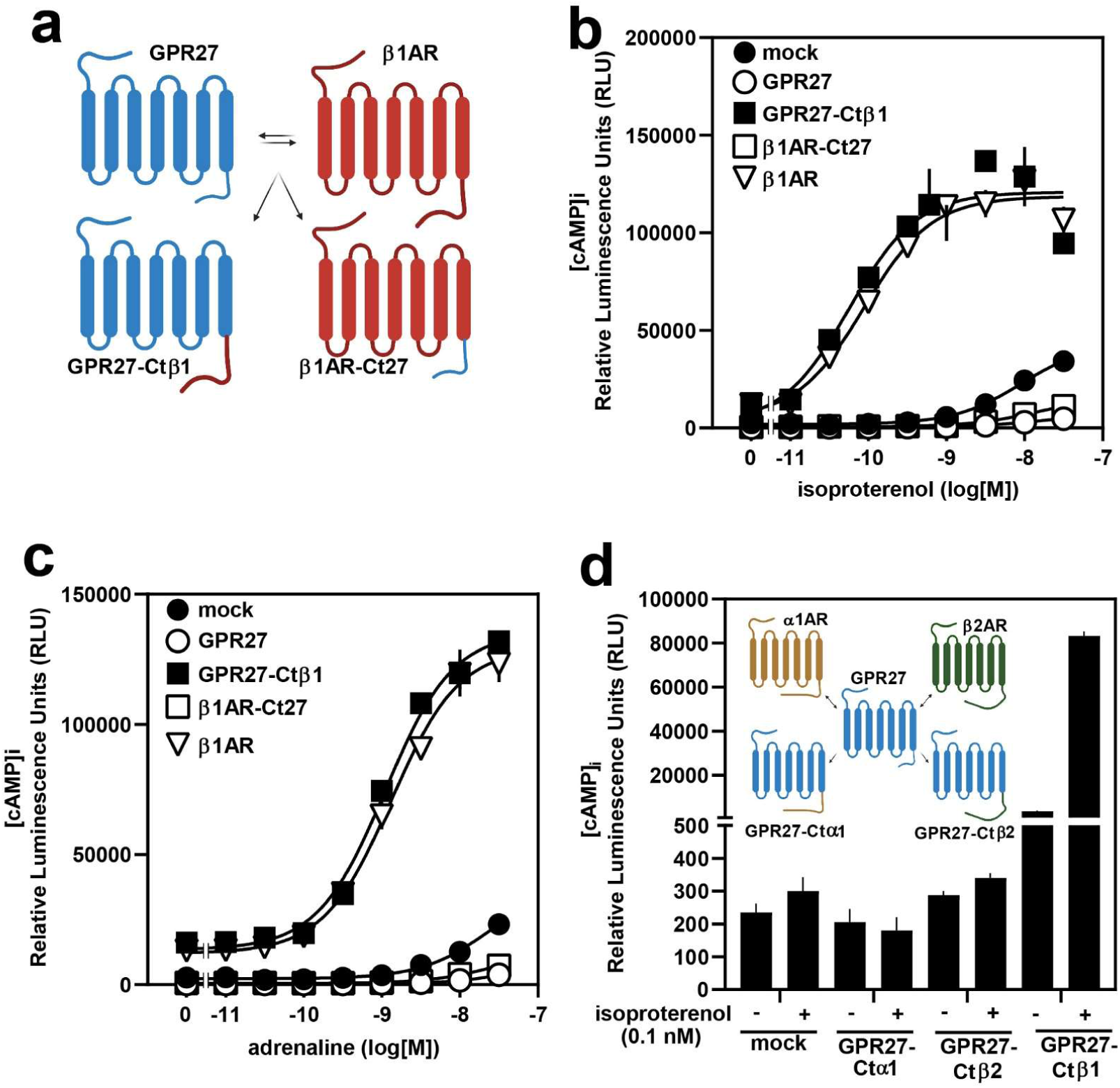
The critical role of β1AR- and GPR27-C-termini in cellular signaling. **(a)** picture illustrating the chimeras between β1AR and GPR27, after the C-termini switch, resulting in GPR27 with the C-terminus of β1AR (GPR27-Ctβ1) and β1AR with the C-terminus of GPR27 (β1AR-Ct27). (b-c) effect of increasing concentrations of isoproterenol (b) and adrenaline (c) on intracellular cAMP levels in HEK293T cells expressing the indicated receptors together with a cAMP-sensitive bioluminescence probe (as described in Material and Methods). (d) (inset) picture depicting the chimeras between GPR27, α1AR and β2AR, resulting in GPR27 with the C-termini of α1AR (GPR27-Ctα1) and of β2AR (GPR27-Ctβ2). Effect of isoproterenol (0.1 nM) on cAMP levels in HEK293T expressing the indicated receptors. Shown are mean values ± SEM, n = 5.

**Table 1.**
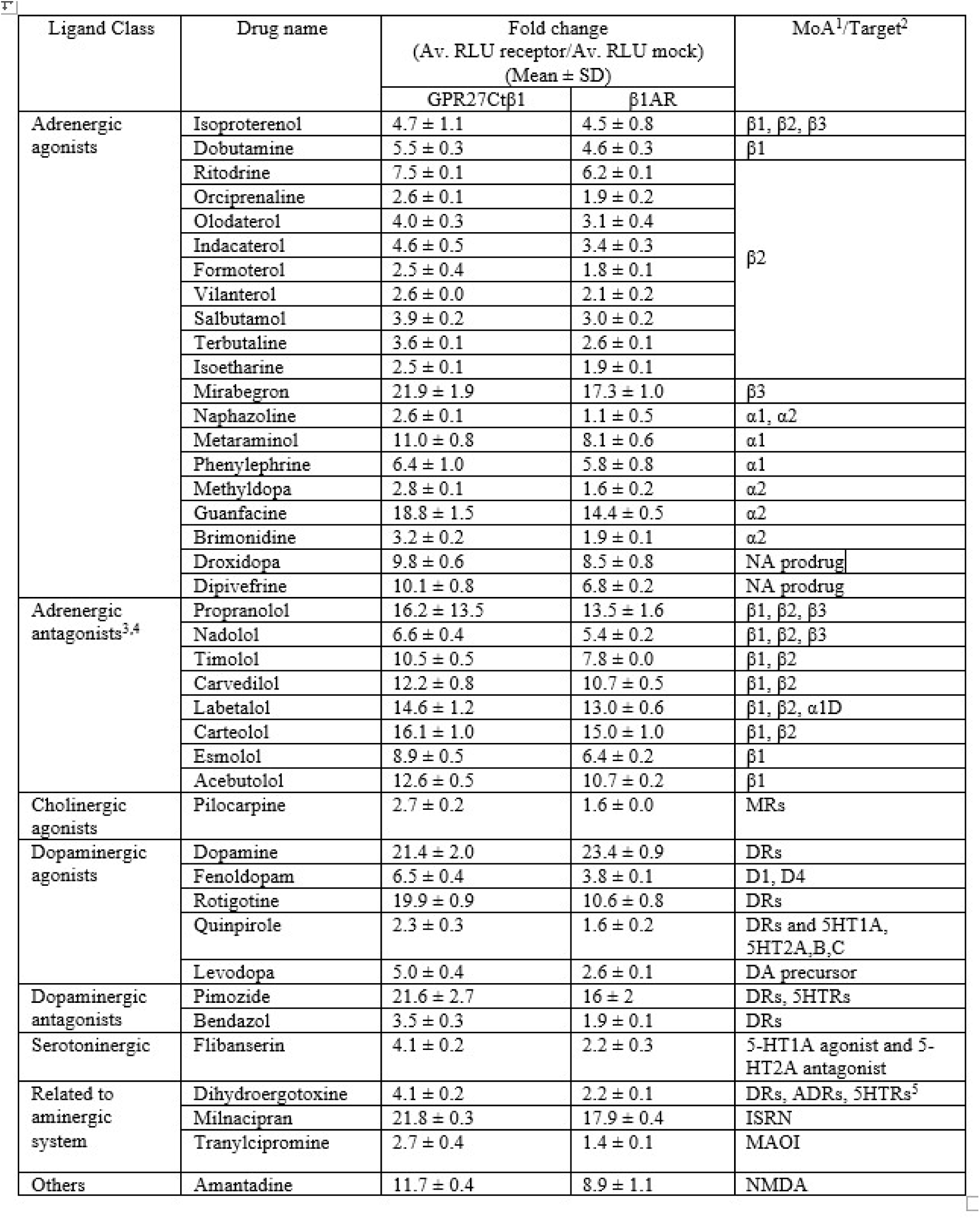
Effect of the shown ligands at 10 µM on intracellular cAMP levels in HEK293T cells expressing GPR27-Ctβ1 chimera and β1AR together with pGlo-22F cAMP sensor. Numbers represent the average of RLU recorded after compound stimulation of GPR27Ctβ1 expressing cells divided by the average of RLU determined after compound stimulation of mock transfected cells.

| Ligand Class | Drug name | Fold change<br>(Av. RLU receptor/Av. RLU mock)<br>(Mean ± SD) | | MoA<sup>1</sup>/Target<sup>2</sup> |
| --- | --- | --- | --- | --- |
| | | GPR27Ctβ1 | β1AR | |
| Adrenergic agonists | Isoproterenol | 4.7 ± 1.1 | 4.5 ± 0.8 | β1, β2, β3 |
| | Dobutamine | 5.5 ± 0.3 | 4.6 ± 0.3 | β1 |
| | Ritodrine | 7.5 ± 0.1 | 6.2 ± 0.1 | β2 |
| | Orciprenaline | 2.6 ± 0.1 | 1.9 ± 0.2 | |
| | Olodaterol | 4.0 ± 0.3 | 3.1 ± 0.4 | |
| | Indacaterol | 4.6 ± 0.5 | 3.4 ± 0.3 | |
| | Formoterol | 2.5 ± 0.4 | 1.8 ± 0.1 | |
| | Vilanterol | 2.6 ± 0.0 | 2.1 ± 0.2 | |
| | Salbutamol | 3.9 ± 0.2 | 3.0 ± 0.2 | |
| | Terbutaline | 3.6 ± 0.1 | 2.6 ± 0.1 | |
| | Isoetharine | 2.5 ± 0.1 | 1.9 ± 0.1 | |
| | Mirabegron | 21.9 ± 1.9 | 17.3 ± 1.0 | β3 |
| | Naphazoline | 2.6 ± 0.1 | 1.1 ± 0.5 | α1, α2 |
| | Metaraminol | 11.0 ± 0.8 | 8.1 ± 0.6 | α1 |
| | Phenylephrine | 6.4 ± 1.0 | 5.8 ± 0.8 | α1 |
| | Methyldopa | 2.8 ± 0.1 | 1.6 ± 0.2 | α2 |
| | Guanfacine | 18.8 ± 1.5 | 14.4 ± 0.5 | α2 |
| | Brimonidine | 3.2 ± 0.2 | 1.9 ± 0.1 | α2 |
| | Droxidopa | 9.8 ± 0.6 | 8.5 ± 0.8 | NA prodrug |
| | Dipivefrine | 10.1 ± 0.8 | 6.8 ± 0.2 | NA prodrug |
| Adrenergic antagonists<sup>3,4</sup> | Propranolol | 16.2 ± 13.5 | 13.5 ± 1.6 | β1, β2, β3 |
| | Nadolol | 6.6 ± 0.4 | 5.4 ± 0.2 | β1, β2, β3 |
| | Timolol | 10.5 ± 0.5 | 7.8 ± 0.0 | β1, β2 |
| | Carvedilol | 12.2 ± 0.8 | 10.7 ± 0.5 | β1, β2 |
| | Labetalol | 14.6 ± 1.2 | 13.0 ± 0.6 | β1, β2, α1D |
| | Carteolol | 16.1 ± 1.0 | 15.0 ± 1.0 | β1, β2 |
| | Esmolol | 8.9 ± 0.5 | 6.4 ± 0.2 | β1 |
| | Acebutolol | 12.6 ± 0.5 | 10.7 ± 0.2 | β1 |
| Cholinergic agonists | Pilocarpine | 2.7 ± 0.2 | 1.6 ± 0.0 | MRs |
| Dopaminergic agonists | Dopamine | 21.4 ± 2.0 | 23.4 ± 0.9 | DRs |
| | Fenoldopam | 6.5 ± 0.4 | 3.8 ± 0.1 | D1, D4 |
| | Rotigotine | 19.9 ± 0.9 | 10.6 ± 0.8 | DRs |
| | Quinpirole | 2.3 ± 0.3 | 1.6 ± 0.2 | DRs and 5HT1A, 5HT2A,B,C |
| | Levodopa | 5.0 ± 0.4 | 2.6 ± 0.1 | DA precursor |
| Dopaminergic antagonists | Pimozide | 21.6 ± 2.7 | 16 ± 2 | DRs, 5HTRs |
| | Bendazol | 3.5 ± 0.3 | 1.9 ± 0.1 | DRs |
| Serotonergic | Flibanserin | 4.1 ± 0.2 | 2.2 ± 0.3 | 5-HT1A agonist and 5-HT2A antagonist |
| Related to aminergic system | Dihydroergotoxine | 4.1 ± 0.2 | 2.2 ± 0.1 | DRs, ADRs, 5HTRs<sup>5</sup> |
| | Milnacipran | 21.8 ± 0.3 | 17.9 ± 0.4 | ISRN |
| | Tranylcipromine | 2.7 ± 0.4 | 1.4 ± 0.1 | MAOI |
| Others | Amantadine | 11.7 ± 0.4 | 8.9 ± 1.1 | NMDA |

### 4.3 GPR27 mediates the inhibitory effects of isoproterenol on transcription factor activity

Based on the observation that the GPR27-Ctβ1 chimera can respond to isoproterenol similarly to β1AR we assumed that GPR27 has the structural requirements to sense adrenergic ligands but it lacks the capacity to signal through G-proteins. To test this idea, we investigated whether GPR27 could affect cellular signaling pathways functions through a more integrative approach, by analyzing the activity of transcription factors in response to isoproterenol. Consistent with the lack of impact on Gs-mediated signaling, stimulation of HEK293T cells expressing GPR27 did not affect the activity of the cyclic AMP response element (CRE)-luciferase (CRE-Luc) fusion protein (Fig. 3a). However, when cells were coexpressed with the serum-response element (SRE)-luciferase (SRE-Luc) fusion protein reporter, isoproterenol stimulation resulted in a GPR27-dependent decrease in SRE-Luc activity. In contrast, β1AR-expressing cells showed the opposite response, with isoproterenol increasing SRE-Luc activity (Fig. 3b). The inhibitory effect of isoproterenol on SRE-Luc activity via GPR27 was concentration-dependent (Fig. 3c) and was completely reversed by the β1AR antagonist metoprolol and the β1/β2AR antagonist propranolol (Fig. 3d). Interestingly, unlike GPR27, the related receptors GPR85 and GPR173 did not influence SRE-Luc activity in response to isoproterenol or other adrenergic ligands (Suppl. Fig. S3). These observations demonstrate that GPR27 specifically inhibits SRE-Luc activity in response to isoproterenol, a distinct feature not observed in GPR85 or GPR173, emphasizing the unique cellular effects mediated by GPR27 within the SREB family.

**Figure 3.**
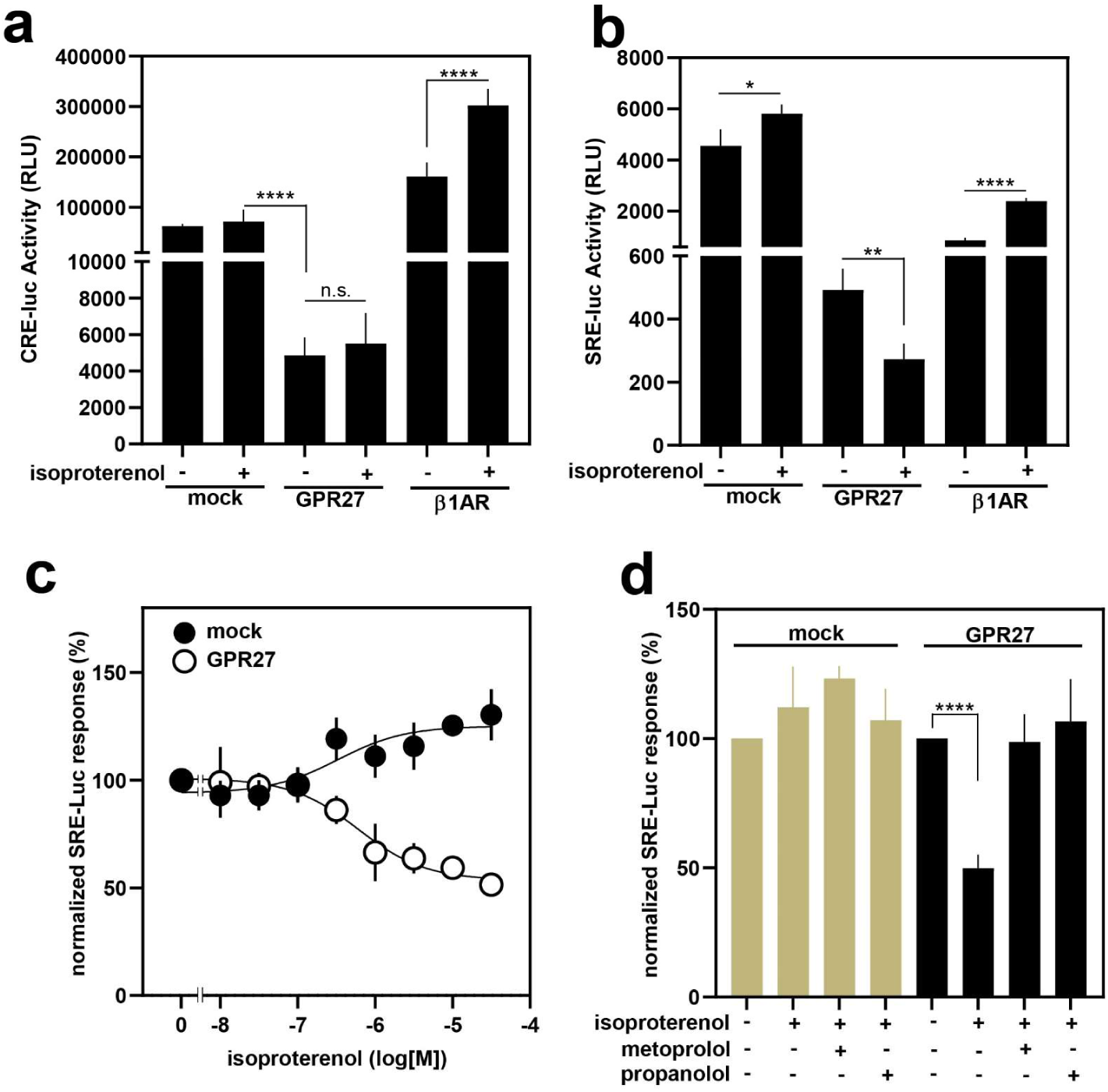
GPR27 mediates isoproterenol-induced inhibition of SRE activity. (a) effect of isoproterenol (1 µM) on the cyclic AMP response element (CRE)-luciferase (CRE-Luc) fusion protein activity in cells expressing empty vector (mock), GPR27, and β1AR. (b) effect of isoproterenol (1 µM) stimulation of HEK293T cells expressing empty vector (mock), GPR27, and β1AR together with the serum-response element (SRE)-luciferase (SRE-Luc) fusion protein reporter. Shown are mean values ± SEM, n = 5. (c) effect of increasing concentrations of isoproterenol on SRE-Luc activity in HEK293T cells expressing empty vector (mock) or GPR27. Data are normalized to the solvent effect on each of the groups. (d) effect of metoprolol and propranolol (10 µM each) on SRE activity in cells expressing empty vector or GPR27 and stimulated with 1 µM isoproterenol. Shown are mean values ± SEM, n = 6, \**P* ≤ 0.05, \*\**P* ≤ 0.001, \*\*\*\**P* < 0.0001, n.s., not significant, (Student’s two-tailed *t*-test). Raw data were normalized to the effect of solvent on each of the samples.

### 4.4 GPR27 mediates the inhibitory effects of isoproterenol on EGF-induced increases in SRE-Luc activity

Previously, we demonstrated that although isoproterenol stimulation of cells expressing GPR27 did not lead to the engagement of Gs- or Gi-protein-mediated signaling pathways, it inhibited SRE-Luc activity. The assay conditions included HEK293T cells expressing GPR27 or empty vector (mock) bathed in DMEM medium supplemented with 2% FBS. This is the reason we hypothesized that the observed inhibition of SRE-Luc activity by GPR27 in response to isoproterenol might be due to the alteration of cellular effects induced by growth factors, such as the epidermal growth factor (EGF), a well-known activator of SRE activity through the EGF receptor (EGFR). As shown in Figure 4a, stimulation of HEK293T cells expressing empty vector (mock) with EGF resulted in a robust increase of SRE-Luc activity. Interestingly, in cells expressing GPR27, the EGF-induced increase of SRE-Luc activity was similar to mock-transfected cells, indicating that the EGF signaling pathway was not affected by the expression of GPR27, as it was in the case of the cAMP pathway (Fig. 1a). However, in cells expressing GPR27, isoproterenol strongly reduced the EGF-induced increase of SRE-Luc activity, an effect not observed in mock-transfected cells (Fig. 4a). These results demonstrate that GPR27 can mediate the inhibition of EGF-induced increase of SRE-Luc activity by isoproterenol, and importantly, in a concentration-dependent manner, with an EC_50_ of around 300 nM (Fig. 4b). To demonstrate the involvement of EGFR in the effects of isoproterenol, cells expressing GPR27 or empty vector were exposed to 1 nM EGF and increasing concentrations of isoproterenol in the presence or absence of EGFR antagonist, erlotinib. As illustrated in Figure 4c, while isoproterenol strongly reduced the effect of EGF on SRE-Luc activity in a concentration-dependent manner only in cells expressing GPR27, erlotinib treatment abolished the concentration-dependency impact of isoproterenol on SRE-Luc activity. Furthermore, in an attempt to determine de structural determinants of GPR27 involved in the inhibition of EGF signaling by isoproterenol, chimeric receptors between GPR27 and β1AR were expressed in HEK293T cells and the effect of isoproterenol stimulation on EGF-induced SRE-Luc activity was determined. When stimulated with increasing concentrations of isoproterenol, GPR27-Ctβ1chimera activated SRE-Luc activity with a similar profile as in β1AR, whereas β1AR-Ct27 responded to isoproterenol in an almost identical manner as GPR27, by concentration-dependently inhibiting the EGF-induced SRE-Luc activation (Fig. 4d). These results correlate with those observed in the functional intracellular cAMP assay (Fig. 2b,c), suggesting that the combination between the GPR27 7TM domains and the C-terminus of β1AR is necessary to obtain a chimera that acquires Gs-protein coupling and SRE-Luc activating characteristics. These evidences point to a critical role of the C-terminus domain in GPR27’s signaling/the signaling of GPR27. To investigate this idea, we generated a truncated version of GPR27 lacking the C-terminal domain (amino acids P359–L375, GPR27ΔCterm). When heterologously expressed in HEK293T cells, GPR27ΔCterm localized at the plasma membrane similarly to GPR27 (Suppl. Fig. S4a), however, contrary to our expectations, showed no coupling to Gs- or Gi-proteins in response to isoproterenol when tested in a functional cAMP assay (Suppl. Fig. S4b,c). Consistent with a lack of Gs- or Gi-coupling, GPR27ΔCterm and β1ARCt27 responded to isoproterenol similarly to GPR27 by inhibiting EGF-induced SRE-Luc activation in a concentration-dependent manner (Fig. 4d). A recent study demonstrated an important role of intracellular loops 3 (ICL3) of β1AR in coupling to Gs proteins (19). To test the potential role of ICL3 of GPR27 and β1AR in receptor functionality, we generated chimera receptors of β1AR having the ICL3 from GPR27 (β1ARicl_3_27) and of GPR27 with the ICL3 from β1AR (GPR27icl_3_β1). Interestingly, β1ARicl_3_27 showed an almost unaffected capacity to respond to isoproterenol in a functional cAMP assay (Suppl. Fig. S5) and SRE-Luc assay, compared to β1AR (Fig. 4e). On the other hand, isoproterenol stimulation of cells expressing GPR27icl3β1 and GPR27icl3β1ΔCt chimeras resulted in a significant reduction of the inhibitory effect of isoproterenol on the EGF-induced increase in SRE-Luc activity (Figure 4e). Interestingly, both chimeras also have a significantly reduced inhibitory effect on the intracellular cAMP levels increased after isoproterenol stimulation compared to cells expressing GPR27 (Fig. 4f), although they showed plasma membrane localization levels similar to GPR27 (Suppl. Fig. S6). These results clearly indicate that the ICL3 of GPR27 appears to be critical in the observed inhibitory effects on SRE-Luc activity and cAMP levels in G-protein independent signaling whereas the C-terminus appears not to be important for these effects.

**Figure 4.**
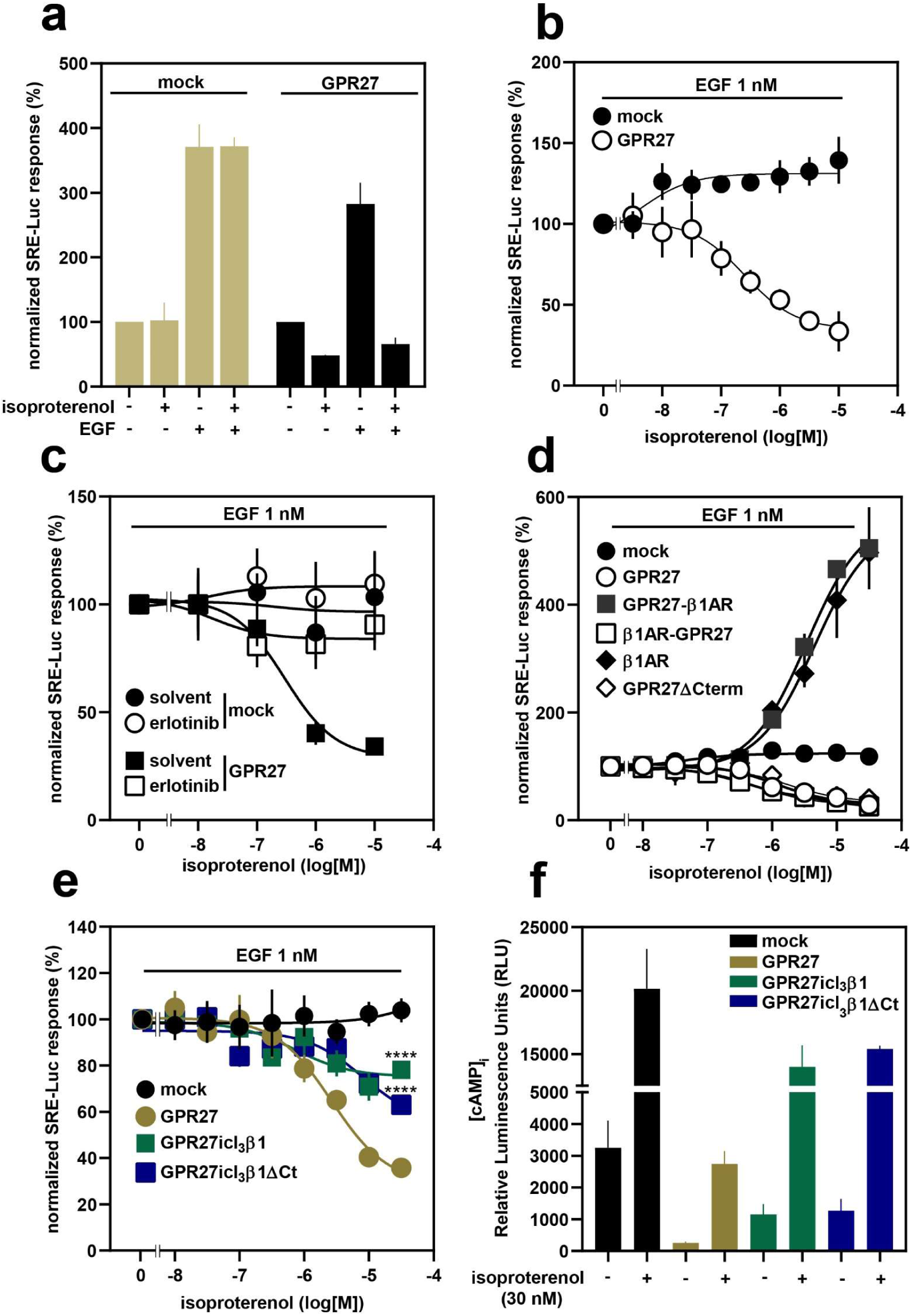
GPR27 mediates the inhibition of isoproterenol on EGF-induced activation of SRE-Luc activity. **(a)** effect of isoproterenol (1µM) on EGF (1nM)-induced SRE-Luc activation in cells expressing empty vector and GPR27. (b) effect of increasing concentrations of isoproterenol on EGF-induced activation of SRE-Luc activity in cells expressing empty vector and GPR27. (c) effect of EGFR antagonist erlotinib (10 µM) on the capacity of increasing concentrations of isoproterenol to inhibit EGF-induced activation of SRE-Luc. (d-e) determination of SRE-Luc activity in cells expressing the indicated receptors and exposed to increasing concentrations of isoproterenol in the presence of 1 nM EGF. (f) determination of intracellular cAMP levels in cells expressing the indicated receptors together with the intracellular cAMP sensitive probe pGlo-22F (Promega) and exposed to 30 nM of isoproterenol. Shown are mean values ± SEM, n = 6, \*\*\*\**P* < 0.0001 (compared with the effect of 30 nM isoproterenol on GPR27 expressing cells) (Student’s two-tailed *t-*test). Raw data were normalized to the effect of solvent on each of the samples.

### 4.5 Isoproterenol-induced inhibition of EGF signaling through GPR27 does not involve G-proteins or arrestins

Based on the data showing that the ligand-induced inhibition of EFG signaling through GPR27 results in a reduced SRE-Luc activity, we wanted to address the natural question: what are the cellular components involved in this phenomenon? To test the potential involvement of G-proteins we assessed the result of GPR27 stimulation with isoproterenol on overall cellular GTPase activity. As illustrated in Suppl. Fig. S7, isoproterenol stimulation of cells expressing β1AR resulted in a significant increase in cellular GTPase activity, as expected. In contrast, exposure to a maximal concentration of isoproterenol did not affect GTPase activity in cells expressing GPR27. To further analyze the potential involvement of specific G-proteins in the observed effects of isoproterenol through GPR27 we employed pharmacological inhibition of Gq/11, G12/13, Gi/o proteins, and the nonselective G-proteins inhibitor, suramin. As shown in Figure 5a, pretreatment of HEK293T cells expressing GPR27 with PTX (Gi/o inhibitor), Y27632 (an inhibitor that prevents G12/13-mediated ROCK-dependent downstream cellular effects) and YM254890 (a Gq/11 inhibitor) did not significantly reverse the inhibitory effect of isoproterenol on EGF-induced increase in SRE-Luc activity. Furthermore, the pretreatment with gallein (an inhibitor of G-protein βγ subunit-dependent signaling) did not reverse the inhibitory effect of GPR27 stimulation with isoproterenol on EGF-induced SRE activation. These results demonstrate that GPR27 mediates the inhibitory effects of isoproterenol on EGF-induced SRE-Luc activation independently of G-proteins, irrespective of their subtype. In response to agonist stimulation GPCRs become substrates for the action of G-protein receptor kinases (GRKs) which are serine-threonine kinases that phosphorylate serine and threonine residues located especially within the ICL3 and C-terminus domain of the activated receptor (20). This phosphorylation event is part of the desensitization mechanism involving arrestin recruitment to the phosphorylated receptor and its internalization (21). It is now well established that G-protein independent signaling of GPCRs, especially through β-arrestins, plays a significant role in regulating cellular responses (22, 23). Since we did not observe any role of G-proteins in the inhibition of SRE-Luc activity by GPR27, we wondered whether GRKs and arrestins could mediate the effects of GPR27 stimulation with isoproterenol on EFG-induced SRE-Luc activation. To test this idea, we first examined the impact of the pharmacological inhibition of GRKs on GPR27-induced inhibition of SRE-Luc activity after isoproterenol stimulation. As shown in Figure 5b, pretreatment of cells expressing GPR27 with the non-selective GRK inhibitor AB06102, could not reverse the inhibition of EGF-induced increase in SRE-Luc activity triggered by stimulation of GPR27 with a maximal concentration of isoproterenol. However, in cells expressing β1AR, pretreatment with AB06102 led to an enhanced SRE activity in response to isoproterenol, as expected considering that the desensitization mechanism of β1AR is affected by the GRKs inhibitor. Furthermore, to explore the possibility that arrestin 2 might mediate the effects of isoproterenol stimulation of GPR27 on SRE activity, we employed a previously described complementation-based system (9), consisting of a chimera GPR27 receptor that has the C-terminus domain replaced with the corresponding domain of the vasopressin 2 receptor (V2R) and fused with a C-terminal portion of the firefly luciferase (Fluc) separated by a linker (GPR27-V2RFluc). We also employed a chimera GPR27 with its C-terminus fused with the C-terminal part of the Fluc (GPR27-Fluc). The β-arrestin 2 fused with the N-terminal part of the Fluc (β-arr-Fluc) was coexpressed with GPR27-V2RFluc and GPR27-Fluc to determine the potential arrestin recruitment to the receptors. As shown in Figure 5c, isoproterenol stimulation of HEK293T cells expressing β-arr-Fluc together with GPR27-V2RFluc or GPR27-FLuc did not increase luciferase activity compared to solvent stimulation, indicating that the complementation of the two fragments of the luciferase was not achieved through isoproterenol stimulated GPR27-V2RFluc. Thus, these results suggest that the arrestin 2 was not recruited to the stimulated chimeric receptor. By contrast, isoproterenol stimulation of chimeric β2AR C-terminally fused to the C-terminal part of the firefly luciferase (β2AR-Fluc) as well as of chimeric β2AR with the C-terminus of V2R fused to the C-terminal part of the firefly luciferase (β2AR-V2RFluc) coexpressed with β-arr-Fluc, resulted in a significant increase in luminescence, demonstrating robust recruitment of β-arr-Fluc to the modified adrenergic receptors following isoproterenol stimulation (Fig. 5c). To verify that the chimeric receptors described above retained their functionality, GPR27-Fluc, GPR27-V2RFluc, β2AR-Fluc and β2AR-V2RFluc were tested for their responsiveness to isoproterenol challenge. As shown in Figure 5d, stimulation with a maximal concentration of isoproterenol led to a strong inhibition of EFG-induced increase in SRE-Luc activity, in the case of cells expressing GPR27-Fluc and GPR27-V2RFluc. In contrast, exposure of cells expressing β2AR-Fluc and β2AR-V2RFluc to a maximal concentration of isoproterenol led to a robust increase in SRE-Luc activity, demonstrating that the modification operated on the receptors did not alter their functionality. All these results indicate that GPR27-mediated inhibition of isoproterenol on the EGF-induced increase in SRE-Luc activity does not involve G-proteins or β-arrestins.

**Figure 5.**
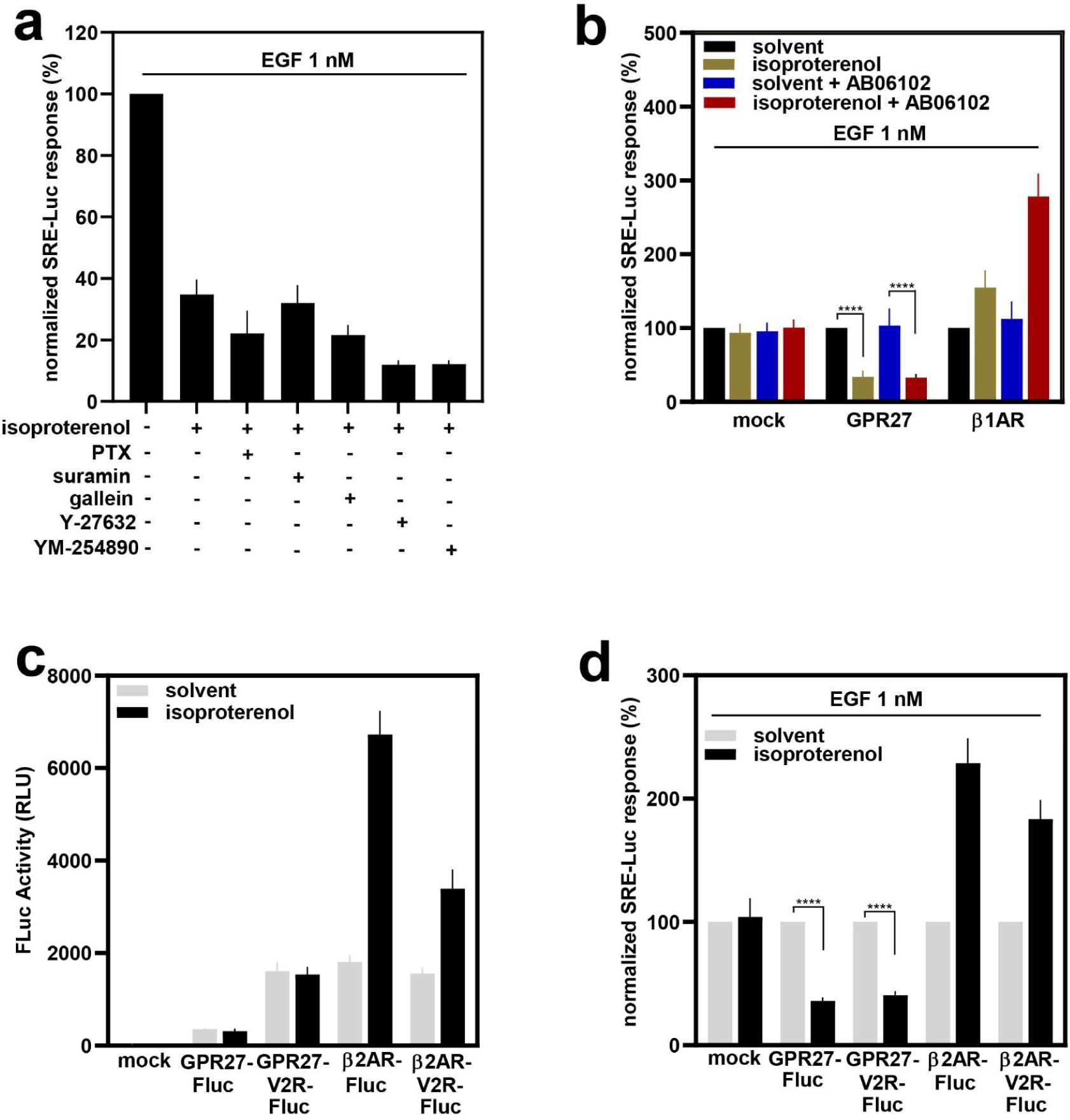
GPR27 mediates the inhibition of isoproterenol on EGF-induced SRE-Luc activation independently of G-proteins and β-arrestin 2. (a) effects of PTX (100 ng/ml), suramin (300 µM), gallein (100 µM), Y-27632 (10 µM) and YM-254890 (100 nM) on isoproterenol-induced inhibition of SRE-Luc activity in the presence of 1 nM EGF in HEK293T cells expressing GPR27. (b) determination of SRE-Luc activity in cells expressing the indicated receptors stimulated with 1 µM isoproterenol, exposed to the GRKs inhibitor AB06102 in the presence of 1 nM EGF. (c) Protein complementation assay in HEK293T cells expressing β-arrestin 2 fused with the N-terminal part of the firefly luciferase (β-arr-Fluc) together with receptors having the C-terminal domain fused to the C-terminal domain of the firefly luciferase (GPR27-Fluc and β2AR-Fluc) or replaced by the C-terminus of vasopressin 2 receptor (V2R) fused to the C-terminus of the firefly luciferase (GPR27V2R-Fluc and β2ARV2-Fluc). The degree of the reconstitution of the luciferase was determined after stimulation of cells with 1 µM isoproterenol and expressed as relative luminescence units (RLU). (d) effect of isoproterenol (1 µM) on EGF-induced SRE activation in HEK293T cells expressing the receptors described in (c). Shown are mean values ± SEM, n = 6, \*\*\*\**P* < 0.0001, (Student’s two-tailed t-test). Raw data were normalized to the effect of solvent on each of the samples.

### 4.6 Isoproterenol-induced inhibition of SRE activity through GPR27 involves dephosphorylation of c-Src and EGFR

In an attempt to identify the intracellular mechanism of SRE inhibition by isoproterenol through GPR27 and based in the observation that this inhibition does not involve G-proteins, we analyzed the involvement of well-known intracellular proteins in the observed effects. Extracellular signal-regulated protein kinase (ERK) 1&2 proteins are well-known modulators of SRE activity (24, 25), therefore it was a natural question whether GPR27 might alter the phosphorylation status of ERK1&2 proteins in response to isoproterenol that would lead to SRE-Luc inhibition. As presented in Figure 6b, stimulation of cells expressing GPR27 with a maximal concentration of isoproterenol did not change the phosphorylation levels of ERK1&2 proteins. However, in response to an equivalent concentration of isoproterenol, β1AR mediated a significant increase in the phosphorylated ERK1&2. Moreover, following isoproterenol stimulation, the GPR27-Ctβ1 chimera mediated a strong increase in the phosphorylation of ERK1&2 proteins, consistent with previously described effects on cAMP levels (Figure 2b). To our surprise, β1ARCt27 chimera stimulation with isoproterenol also led to significant phosphorylation of ERK1&2 (Fig. 6c), although it mediated inhibition of EGF-induced SRE activation, in a similar manner as GPR27 (Fig. 4c). These results indicate that the inhibition of SRE activity by GPR27 in response to isoproterenol does not occur through ERK1&2 proteins. Since the observed cellular effects of GPR27 in response to isoproterenol are the inhibition of EGF-induced SRE activation, we examined whether there is an effect of GPR27 on EGFR phosphorylation status or a direct interaction between the two proteins. Although we could not see a ligand-induced interaction between GPR27 and EGFR, as revealed by immunoprecipitation studies (Suppl. Fig. S8), isoproterenol stimulation of cells expressing GPR27 led to a significant reduction of EGF-induced phosphorylation level of EGFR (Fig. 6d).

**Figure 6.**
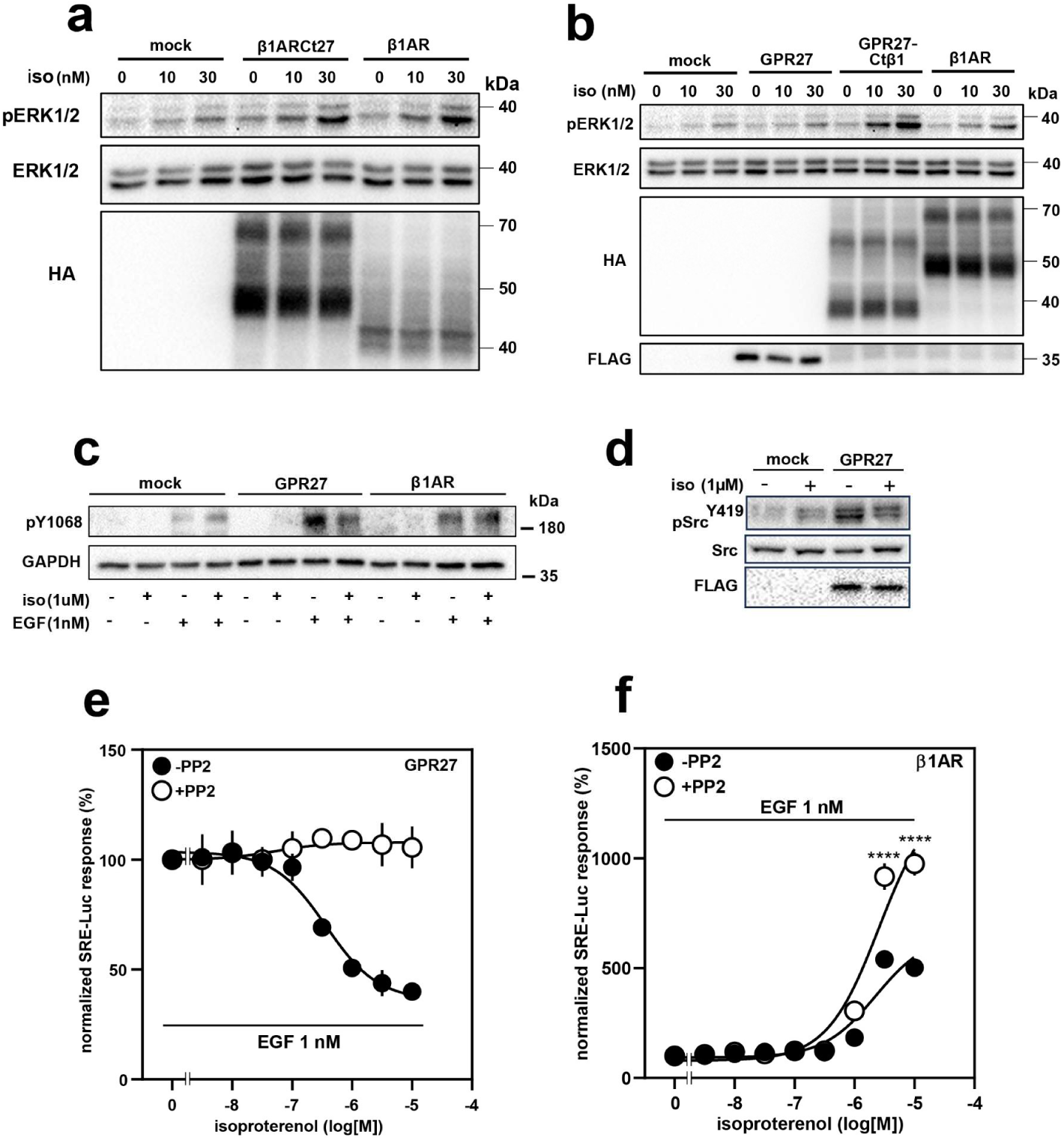
Isoproterenol inhibition of EGF-induced SRE activation through GPR27 involves dephosphorylation of EGFR and s-Src, independently of the MAP kinase pathway component, ERK1&2. (a-b) phosphorylation of ERK1/2 proteins in response to isoproterenol stimulation of HEK293T cells expressing the indicated receptors as revealed by immunoblotting using anti-phospho-p44/42 MAPK (Erk1/2) antibody, (pERK1/2, upper blot). Loading controls for each sample and expression of the indicated receptors were determined by immunoblotting the same membrane with ERK1/2 and anti-HA (a) and anti-FLAG (b) antibodies. (c) The phosphorylation of endogenous EGFR in response to isoproterenol stimulation of HEK293T cells expressing the indicated receptors was revealed by immunoblotting using an anti-phospho-EGF Receptor (pY1068 on the presented blot) antibody. Loading control of each sample was determined by immunoblotting using an anti-GAPDH antibody. (d) effect of isoproterenol stimulation of HEK293T cells expressing GPR27 and empty vector, on the phosphorylation levels of endogenous c-Src as revealed by immunoblotting with anti-phospho-c-Src antibody (pSrc^419^ on the presented blot). Loading control levels and expression of GPR27 were visualized using anti-Src and anti-FLAG antibodies. (e-f) effect of Src family inhibitor, PP2, on isoproterenol-induced inhibition of SRE activity in the presence of EGF (1 nM) in HEK293T cells expressing GPR27 (e) and β1-AR (f). Shown are mean values ± SEM, n = 6, \*\*\*\**P* < 0.0001, (Student’s two-tailed t-test).

The role of c-Src proteins in phosphorylation of EGFR (26) as well as in EGFR transactivation by GPCRs has been previously demonstrated (27). Therefore, an obvious question was whether isoproterenol-stimulated GPR27 affects the phosphorylation levels of c-Src. As illustrated in Figure 6e, stimulation of cells expressing GPR27 led to dephosphorylation of c-Src, a result in good agreement with that observed in the case of EGFR phosphorylation. Interestingly, cells expressing GPR27 have an increased basal (in the absence of isoproterenol stimulation) level of the phosphorylated c-Src compared to control transfected cells. Consistent with the role of Src in mediating transinhibition of EGFR by isoproterenol through GPR27, pretreatment of cells expressing GPR27 with the c-Src inhibitor PP2, led to a complete loss of the inhibitory effect of isoproterenol on EGF-induced SRE activation (Fig. 6f). Interestingly, in cells expressing β1AR, pretreatment with PP led to a potentiation effect of isoproterenol-induced SRE activation (Fig. 6g). Based on these results we concluded that GPR27 mediates the inhibition of EGF-dependent activation of SRE through a previously undescribed transinhibition phenomenon, involving dephosphorylation of c-Src and EGFR.

## 5. Discussion

In the present study, we demonstrate that GPR27 is a receptor that transinhibits RTKs such as EGFR in response to adrenergic ligands. Our starting point was the search for cellular effects induced by the heterologous expression of SREBs (GPR27, GPR85, and GPR173). Interestingly, GPR27 but not GPR85 and to a lesser extent GPR173 induced significant inhibition of the adenylyl-cyclase/cAMP pathway in a Gi-proteins-independent manner. Our hypothesis that the C-terminus domain of GPR27 might play a role in the observed cellular effects of GPR27 led us to the generation of a series of chimeric receptors consisting of GPR27 lacking the C-terminus, and with the C-terminus replaced by the homologous domain of GPCRs with known ligands. Conversely, we generated chimeras consisting of known GPCRs with their C-terminus replaced by the homologous domain of GPR27. All these chimeras were analyzed for expression and localization at the plasma membrane followed by functional analyses. Surprisingly, among all the properly expressed chimeras, GPR27-Ctβ1 functioned similarly to β1AR regarding Gs-proteins coupling and responsiveness to adrenergic agonists and antagonists. On the other hand, β1AR-Ct27 lost its ability to couple to Gs proteins in response to agonists, suggesting that the C-terminus of GPR27 severely impacts chimera signaling through Gs-proteins. These two important pieces of evidence prompted us to generate and test the functionality of GPR27ΔCt. Contrary to our expectations, the C-terminally truncated receptor did not regain G-proteins-coupling in response to adrenergic ligands, indicating that the C-terminus might not be the only domain critically involved in GPR27’s particular signaling characteristics. Although GPR27-Ctβ1 functioned similarly to β1AR, GPR27 did not respond to adrenergic ligands by engaging canonical G-proteins mediated signaling pathways. These results led us to interrogate more integrative signaling pathways such as determining transcriptional factor activity after receptor stimulation. Among all the transcriptional factors tested, only EGF-induced increase in SRE activity was strongly inhibited by GPR27 after stimulation with adrenergic ligands. Interestingly, the observed inhibitory effect of GPR27 was insensitive to G-proteins inhibitors, did not involve arrestin recruitment, and did not alter the levels of the phosphorylated ERK1&2 proteins, suggesting a novel cellular signaling mechanism of GPR27. Importantly, GPR27-Ctβ1 functioned similarly to β1AR regarding SRE-Luc activation in response to isoproterenol whereas β1AR-Ct27 was similar to GPR27, inducing an opposite effect on EGF signaling through SRE. These observations point to a critical role of the C-terminal domain of GPR27, especially when considering its severe impact on β1AR signaling. Unexpectedly to us, GPR27ΔCt still displayed full signaling characteristics of GPR27, also regarding the effects on SRE activity. This is the reason we investigated the potential role of ICL3 of GPR27 and β1AR, by generating chimeras between both receptors on that domain. Notably, β1ARicl_3_27 functioned as β1AR in terms of Gs-proteins coupling and SRE activation, however, GPR27icl_3_β1AR had a significantly lower capacity to inhibit EGF signaling through SRE, illustrating its importance in GPR27’s atypical signaling features. This assertion was supported by the fact that expression of GPR27icl3β1AR strongly reduced the inhibition on isoproterenol-induced cAMP accumulation compared to GPR27 partially answering our starting question derived from the observation of the strong effects of GPR27 on cAMP levels (Fig. 1a). However, GPR27icl_3_β1AR did not regain G-protein engagement capacity in response to isoproterenol and could still inhibit EGF signaling through SRE although with a significantly reduced efficacy. Interestingly, GPR27icl_3_β1ΔCt functioned similarly to GPR27icl_3_β1AR, pointing to a less critical role of the C-terminus in the chimera’s functionality. In an attempt to identify the intracellular signaling pathways involved in the effects of GPR27 on EGF signaling, we examined the tyrosine-phosphorylation levels of proteins known to be involved in the transactivation of EGFR by GPCRs, such as endogenous c-Src as well as at the tyrosine-phosphorylation of EGFR itself. Stimulation of cells expressing GPR27 with isoproterenol led to dephosphorylation of c-Src and EGFR, thus partially explaining the inhibitory effects of GPR27 on SRE activity. The involvement of c-Src in the transinhibition of EGFR by GPR27 was further supported by the effect of c-Src family of tyrosine kinases proteins inhibitor, PP2, which completely abolished the concentration dependency of isoproterenol on SRE-Luc activity in cells expressing GPR27. Intriguingly, PP2 had a rather positive effect on β1AR-induced activation of SRE-Luc activity in response to isoproterenol stimulation, a fact that deserves future attention. An even more interesting observation is the absence of the involvement of ERK1&2 in the inhibition of SRE-Luc activity after isoproterenol stimulation of GPR27.

Based on the results presented in this study, we support the notion that GPR27 senses extracellular adrenergic ligands, and in turn dephosphorylates c-Src and EGFR proteins through an unknown mechanism, but likely independent of G-proteins and β-arrestins. Notably, several intriguing pieces of evidence raise important questions that need to be further addressed. For example, the β1AR-Ct27 chimera which mimics GPR27 in terms of SRE inhibition and the lack of G-protein engagement, increased in fact the phosphorylation levels of ERK1&2 proteins in response to isoproterenol exposure. Also intriguingly, cells expressing GPR27 showed increased phosphorylated levels of c-Src proteins in the absence of isoproterenol stimulation. This effect correlated with the increased levels of phosphorylated EGFR in cells expressing GPR27 and stimulated with EGF, compared to control- and β1AR-transfected cells. This could suggest that GPR27 might have a basal “phosphatase inhibitory”-like effect, which is reversed by stimulation with adrenergic ligands. Importantly, we could neither see a direct interaction between GPR27 and EGFR nor a ligand-induced one, supporting the idea that the transinhibition of EGFR by GPR27 occurs through the engagement of cellular effectors such as Src family of tyrosine kinases proteins. It would be important to clarify whether there is an interaction between GPR27 and the endogenously expressed β1AR that could lead to the formation of a heteromer with a completely different function than a classical Class A GPCR, also an exciting hypothesis since no example of such a heteromer has been so far reported. However, this hypothesis might have its weaknesses based on our data demonstrating the absence of β1- or β2-adrenergic receptors in HEK293T as they do not respond to dobutamin, at concentrations up to 30 µM (Suppl. Fig. 9).

Although GPR27 belongs to the SREB family of receptors, together with GPR85 and GPR173, it distinguishes itself by sharing a similar β1AR-like ligand selectivity but with opposed functionality, features not shared by GPR85 and GPR173. Ligand-dependent transinhibition of RTKs by GPCRs has not been described so far and the cellular mechanism underlying this phenomenon might be complex involving cellular effectors that remain to be yet characterized in further studies. However, published studies on the potential role of GPR27 reveal antitumor effects (10) that could be explained by our results on the transinhibition of RTKs by GPR27. In conclusion, we provide evidence for a new cellular phenomenon, transinhibition of EGFR by the orphan GPR27, which appears to be a novel and atypical adrenergic receptor.

