## Supplementary material for "GPR27 mediates adrenergic ligands-induced transinhibition of EGFR": Suppl Figures

### Supplementary Figures Legends

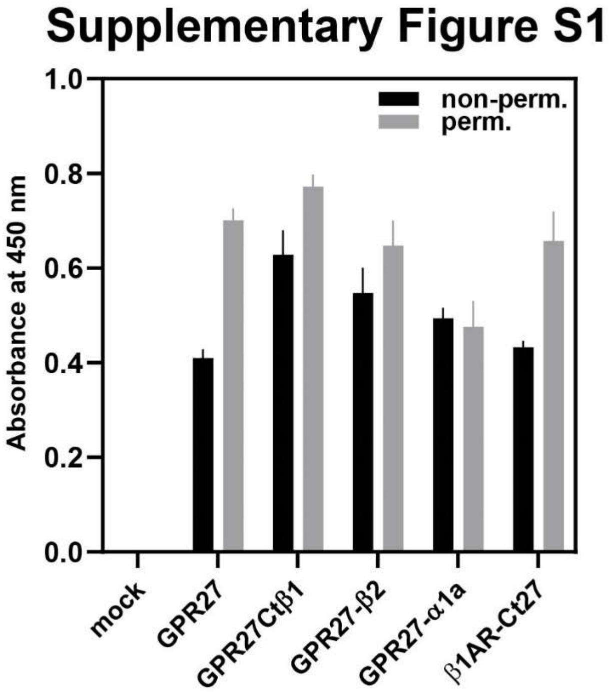


**Suppl. Fig. S1**

Whole-cell ELISA on HEK293T cells expressing the indicated receptors in non-permeabilized conditions (non-perm) to show plasma membrane localization or in permeabilized conditions (perm.) to determine the overall expression of the indicated receptors, using an anti-HA-HRP-conjugated antibody. Shown are mean values ± s.e.m, n = 5.

### Supplementary Figure S2

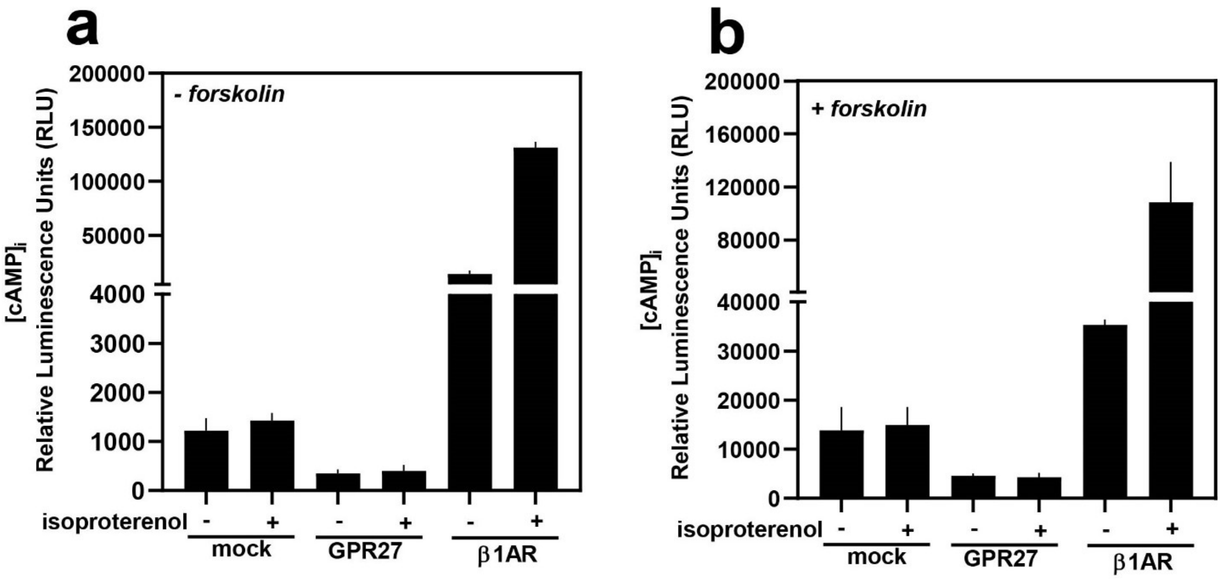


**Suppl. Fig. S2**

(a-b) Determination of intracellular cAMP levels in HEK293T cells expressing the indicated receptors together with the cAMP-sensitive probe after stimulation with 1 nM isoproterenol in the absence (a) or presence of 10 µM isoproterenol (b). Shown are mean values ± s.e.m, n = 5.

### Supplementary Figure S3

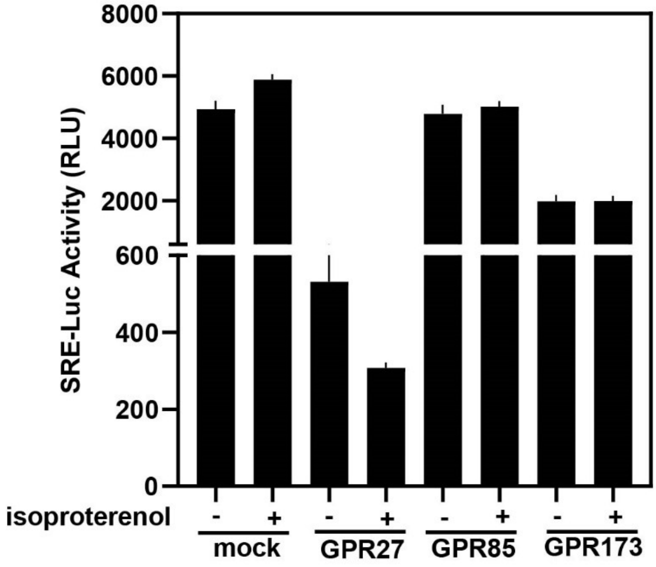

**Suppl. Fig. S3**

Effect of 1 μM isoproterenol on SRE-Luc activity in HEK293T cells expressing the indicated receptors. Shown are mean values ± s.e.m, n = 5.

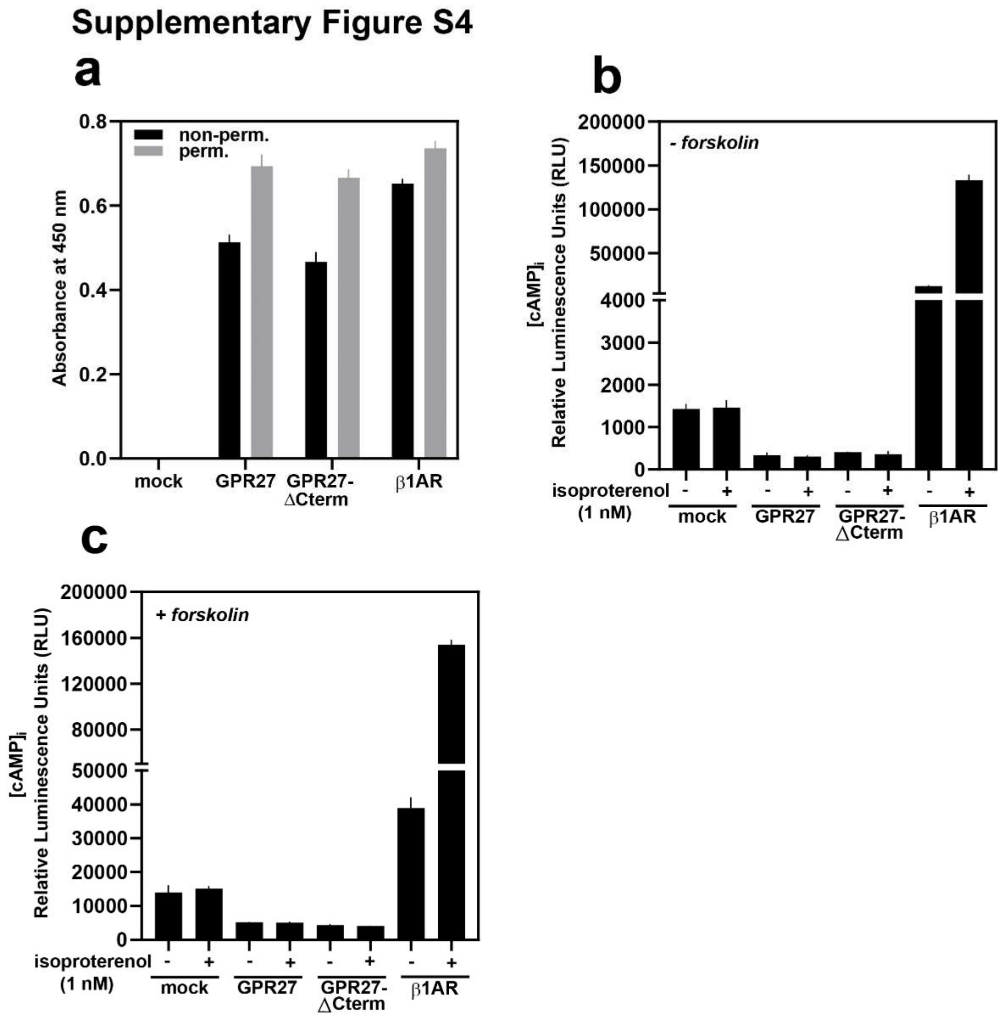


**Suppl. Fig. S4**

(a) Whole-cell ELISA on HEK293T cells expressing the indicated receptors to determine plasma membrane localization (non-perm.) and overall cellular expression (perm.). (b-c) Determination of intracellular levels of cAMP in HEK293T cells expressing the indicated receptors together with the cytosolic localized cAMP-sensitive probe after stimulation with 1 nM isoproterenol, in the presence (b) or absence (c) of forskolin (10 μM). Shown are mean values ± s.e.m, n = 5.

### Supplementary Figure S5

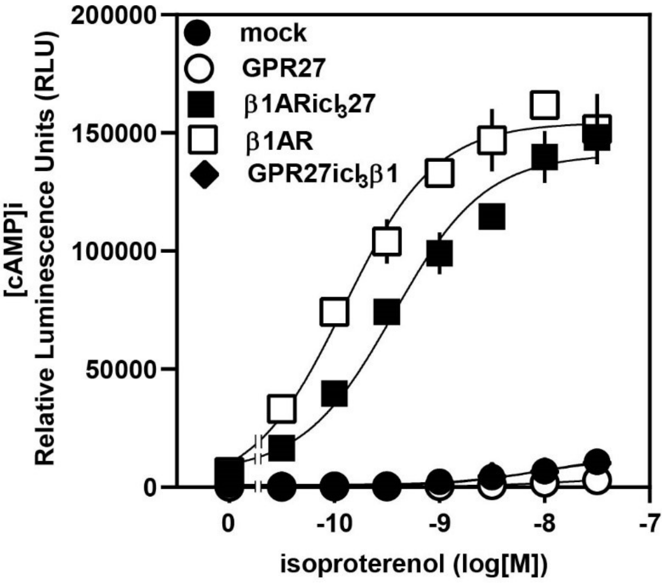


**Suppl. Fig. S5**

Effect of increasing concentrations of isoproterenol on intracellular levels of cAMP in HEK293T cells expressing the indicated receptors together with the cytosolic localized cAMP sensitive probe. Shown are mean values ± s.e.m, n = 5.

#### Supplementary Figure S6

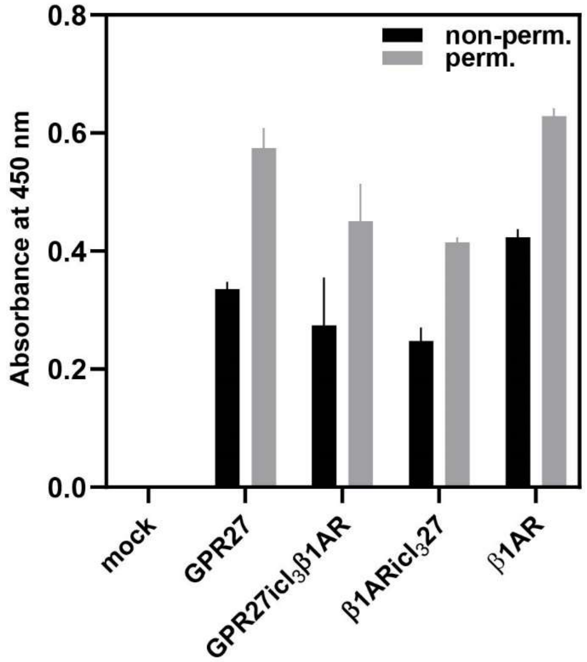


**Suppl. Fig. S6**

Whole-cell ELISA on HEK293T cells expressing the indicated receptors to evaluate plasma membrane localization (non-perm) and total cellular expression (perm.) using anti-HA-HRP conjugated antibody. Shown are mean values ± s.e.m, n = 5.

#### Supplementary Figure S7

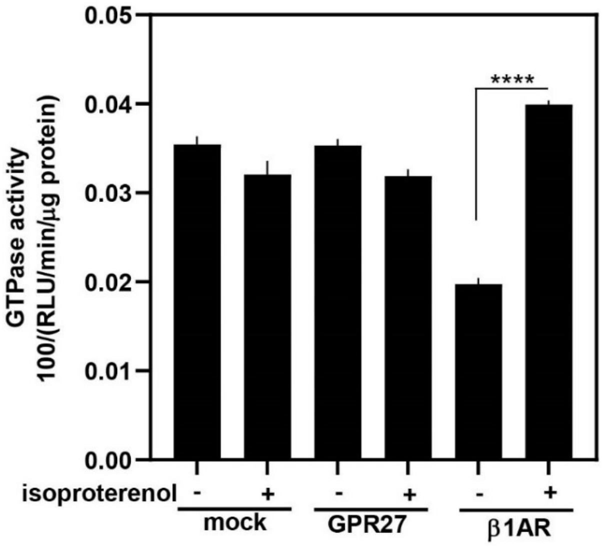

**Suppl. Fig. S7**

Total cellular GTPase activity assay in HEK293T cells expressing the indicated receptors performed as described in Materials and Methods. For easier visualization, the raw results were divided to one hundred. Shown are mean values ± s.e.m, n = 3. ****$P < 0.0001$, (Student's two-tailed *t*-test).

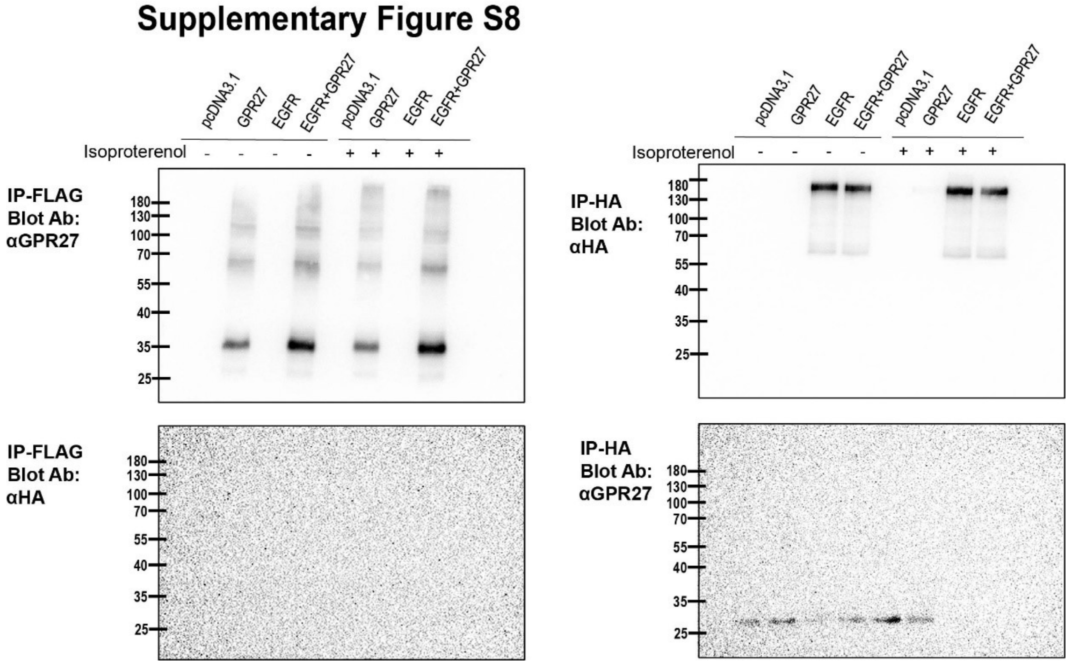


**Suppl. Fig. S8**

Immunoprecipitation followed by Western Blotting of cellular lysates (input: 300 µg total protein) from cells expressing N-terminally FLAG tagged GPR27 and C-terminally HA tagged EGFR. Where indicated, cells were exposed for 10 minutes to 1 µM isoproterenol. IP-FLAG, immunoprecipitation of total cellular lysates using anti-FLAG antibody; IP-HA, immunoprecipitation of total cellular lysates using anti-HA antibody. The assay was performed as presented in the Material and Methods.

### Supplementary Figure S9

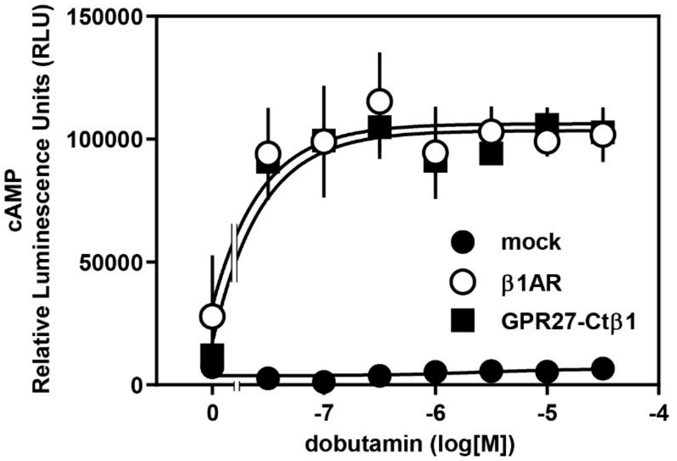


**Suppl. Fig. S9**

Effect of increasing concentrations of dobutamin on intracellular levels of cAMP in HEK293T cells expressing the indicated receptors together with the cytosolic localized cAMP sensitive probe. Shown are mean values ± s.e.m, n = 6.
